# Differences in stomatal conductance between leaf shape genotypes of *Ipomoea hederacea* suggest divergent ecophysiological strategies

**DOI:** 10.1101/2025.02.01.636047

**Authors:** Yash Kumar Singhal, Julia Anne Boyle, John R. Stinchcombe

## Abstract

Intraspecific variation in leaf shape affects many physiological processes but its impact on plant distribution is underexplored. Using a common garden, we studied daytime thermoregulation of lobed and entire leaf genotypes of *Ipomoea hederacea*, which displays a latitudinal leaf shape cline. Both leaf shapes maintained similar temperatures but entire leaf genotypes had significantly increased stomatal conductance in warmer/sunnier weather. With less potential water loss, lobed genotypes may have advantages in drier conditions. Lobed genotypes are more common in the north of the species’ range, which receives less summer precipitation, suggesting water availability as a potential clinally varying selective agent.

## Description

A fundamental goal of ecology and evolution is to understand the distribution of intraspecific variation, as it affects responses to selection within and across environments (DeMarche et al., 2019; Song and Li 2023). Climate change is expected to greatly change selective landscapes (Parmesan and Hanley 2015; Chiang et al., 2021), and plant species are predicted to undergo latitudinal range-shifts to track optimum environmental conditions (Chen et al., 2011; Lenoir and Svenning 2015; Tomiolo and Ward 2018). Species persistence will rely, in part, on existing genetic variation as well as phenotypic and physiological plasticity (Anderson et al., 2012; Bellard et al., 2012). Here, we examine how genetic variation in leaf morphology in *Ipomoea hederacea* (Ivy-leaf morning glory) leads to differences in ecophysiology, and whether these differences are consistent with existing phenotypic clines.

There is an incredible diversity of leaf shapes between and within species, and leaves are involved in crucial processes such as photosynthesis, thermoregulation, and defence (Nicotra et al., 2011). A single Mendelian locus controls leaf shape in *Ipomoea hederacea*, resulting in a homozygous lobed genotype, a homozygous entire genotype, and an intermediately lobed heterozygote (Figure 1a; Elmore 1986; Bright 1998). The species’ range extends from Mexico to southern Pennsylvania with a latitudinal cline in leaf shape, where lobed plants dominate northern populations while southern populations are either entire or polymorphic (Bright 1998; Campitelli and Stinchcombe 2013a). The agents of selection maintaining this cline remain unclear (Campitelli and Stinchcombe 2013b).

**Figure 1.**
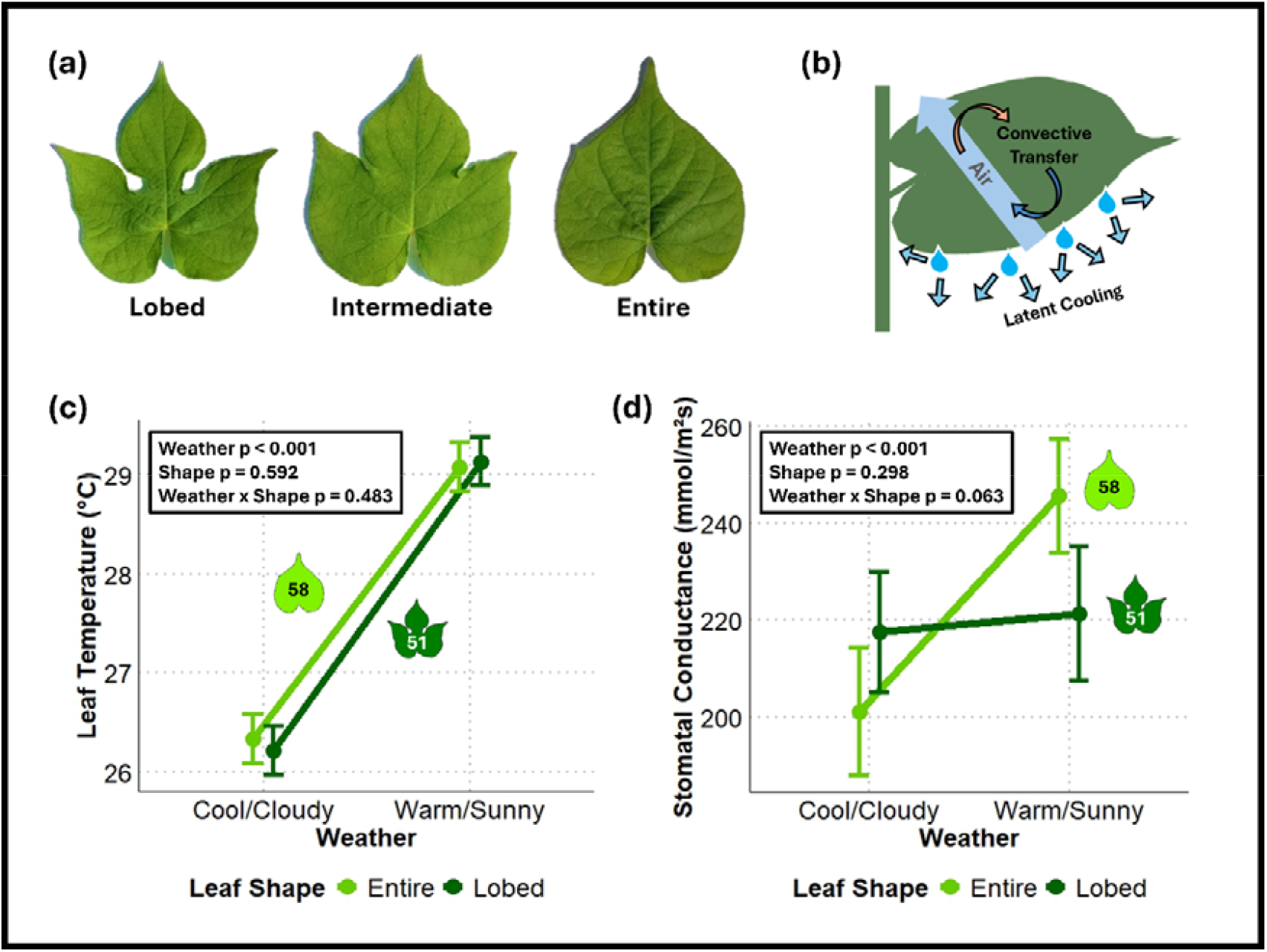
Differing ecophysiology between leaf shape genotypes of *Ipomoea hederacea*: (a) Leaf shape phenotypes of *Ipomoea hederacea*. (b) Schematic showing two major modes of leaf thermoregulation, convective transfer and latent cooling. (c) Daytime leaf temperature varies with abiotic environment but not leaf shape. (d) Stomatal conductance varies with abiotic environment for entire leaf plants but not lobed genotypes. Overall linear model statistics are displayed on the graphs in (c) and (d). Points represent marginal means ± standard error.

Plants thermoregulate through three main mechanisms (Figure 1b): radiative heat transfer (heat exchange in the form of radiation), convective heat transfer (exchange of heat with surrounding air), and latent cooling (loss of heat through transpiration; Nobel 2020). Ecophysiology theory predicts that leaf shape influences thermoregulation as it affects air flow around leaves and thus convective heat transfer between leaf surfaces and surrounding air (Vogel 1970; Schuepp 1993). Lobed leaves are predicted to have smaller effective boundary layers and thus more efficient convective cooling, which keeps them more closely matched to air temperature than entire leaves (Vogel 1970; Gurevitch and Schuepp 1990, Schuepp 1993; Nobel 2020).

We grew *I. hederacea* of all leaf shape genotypes in a long-term common garden at a northern latitude, and measured leaf temperature and abaxial stomatal conductance to characterize thermoregulation. We repeated these measurements on two days differing in temperature and light (Table 1): a cool/cloudy day and a warm/sunny day 48 hours apart, to study the same plants under different environmental conditions.

**Table 1.** Average weather metrics calculated from 181 measurements, taken every minute between 12:00 and 15:00 hrs (on July 24th and July 26th, 2024) at the Koffler Scientific Reserve Weather Station, near the common garden. All weather metrics except precipitation were significantly different between the two days (p < 0.001).

| Day | Average Air Temperature (°C) | Average PAR ($\mu\text{E}$) | Average Pressure (mbar) | Average Wind Speed (m/s) | Average Gust Speed (m/s) | Total Rain (mm) |
| --- | --- | --- | --- | --- | --- | --- |
| July 24, 2024 (cool/cloudy) | $23.8 \pm 0.067$ | $678 \pm 26.1$ | $977 \pm 0.033$ | $1.12 \pm 0.048$ | $2.76 \pm 0.071$ | $0.00 \pm 0.00$ |
| July 26, 2024 (warm/sunny) | $24.9 \pm 0.045$ | $1730 \pm 2.52$ | $985 \pm 0.020$ | $0.897 \pm 0.049$ | $2.17 \pm 0.067$ | $0.00 \pm 0.00$ |

Consistent with previous results (Campitelli et al., 2013; Campitelli and Stinchcombe 2013b), there was no significant effect of leaf shape on daytime leaf temperature on either day (Tukey’s post hoc test, p = 0.595 on the cloudy day and 0.775 on the sunny day); leaves on all plants were on average 2.83°C warmer in warm/sunny conditions (Tukey’s post hoc test, p < 0.001 for both shapes; Figure 1c). Increased leaf temperature was coupled with a significant 44.45 mmol/m^2^s increase in average stomatal conductance of entire shaped leaves in warm/sunny conditions (Tukey’s post hoc test, p = 0.027; Figure 1d), but not lobed leaves (Tukey’s post hoc test, p = 0.856; Figure 1d). Higher entire leaf stomatal conductance suggests increased investment into latent cooling, as plants were losing more water through their stomata when it was hotter. As lobed leaves did not increase latent cooling in hotter temperatures, yet all plants maintained similar leaf temperatures, plants with lobed leaves may thermoregulate via another strategy, potentially through greater convective cooling facilitated by their shape.

Stomatal conductance affects water loss and gas exchange (Cowan 1982) which is strongly correlated with photosynthesis (Lawson et al., 2018). Thermoregulation and photosynthetic activity are thus likely both altered between genotypes. We suggest that reduced dependency on latent cooling may allow lobed genotypes to keep their stomata closed, reduce evaporative water loss, and conserve water on hot and sunny days, which would be beneficial in areas with low water availability. In contrast, greater stomatal conductance in entire genotypes on sunny days implies more open stomata, facilitating greater latent cooling, water loss, and gas exchange. Such a strategy would be advantageous in areas with greater water availability. Our hypothesis is consistent with the leaf shape cline in *I*.

*hederacea*, where lobed genotypes are more frequent in northern latitudes which receive less summer precipitation (e.g., Martin-Benito and Pederson 2015), and entire genotypes are more common in southern sites with greater summer precipitation. These findings suggest water availability as a potential selective agent contributing to the leaf shape cline.

Future work could test the hypothesis of divergent strategies by measuring biomass accumulation, photosynthetic efficiency, and fitness of the genotypes under different water availability conditions in the field. Experiments that factorially manipulate temperature and photosynthetically active radiation (PAR) could also test whether thermoregulatory strategy and photosynthetic activity explain the trends in stomatal conductance. In addition, it would be useful to explicitly measure and compare boundary layer conductance between the two leaf shape genotypes under natural conditions.

Intraspecific variation in thermoregulation and other physiological traits is likely a factor underlying species distributions (Westerband et al., 2021) and may affect species resilience to ecological challenges (Niu et al., 2020). One implication of our results is that species and genotypes that rely strongly on latent cooling or have other water-intensive physiological traits may face novel ecological challenges with changing precipitation patterns.

## Methods

We used an experimental population of *I. hederacea* grown north of their natural range at the Koffler Scientific Reserve (KSR, www.ksr.utoronto.ca) in Ontario, Canada (44°01’48”N, 79°32’01”W). We started by collecting both genotypes from North Carolina, USA, and selfing them for seven generations to create the parents (P1). We crossed homozygous entire and lobed P1 individuals; an F1 heterozygote was selfed to produce an F2 population (Campitelli and Stinchcombe 2013b; Boyle et al., 2024). We allowed heterozygous F2 individuals to self-fertilize, producing F3 seeds planted into the common garden in 2019. We let plants naturally dehisce in 2019 and each subsequent year, ploughing the field in early summer each year to maintain the population. We sampled the 5th generation in July 2024, measuring 58 entire and 51 lobed individuals (heterozygotes and lobed grouped). We grouped heterozygous and homozygous lobed individuals because our ecophysiological hypotheses were based on differences between lobed and entire shaped leaves; distinguishing between heterozygous and homozygous lobed individuals in the field is also challenging. Because heterozygotes are intermediately lobed, considering them with homozygous lobed individuals should make our results conservative.

### Measurements

We measured leaf temperature (°C) and abaxial stomatal conductance (mmol/m^2^s) using a hand-held leaf porometer (Decagon Devices, SC-1 Model). We calibrated and used the device without a desiccant. We surveyed the field systematically using quadrats of equal size, and measured the first fully open leaf from the top of each plant; measured leaves were of similar size. We took measurements of the same plants on a cool/cloudy day (July 24th, 2024) and a warm/sunny day (July 26th, 2024) to study thermoregulation under different environmental conditions (Table 1). All measurements were taken between 12:00 and 15:00 hrs, based on a preliminary time-series.

### Weather

We gathered weather data from a weather station 530 m from the site (Table 1) using measurements from every minute between 12:00 and 15:00 hrs. The cool/cloudy day of sampling (July 24th) had a lower average air temperature, average PAR, average pressure, average wind-speed, and average gust-speed (Welch Two Sample t-test, all p < 0.001) than the warm/sunny day (July 26th). There was no precipitation during the sampling interval on either day.

### Statistical analysis

We analyzed data using R (v4.4.1; R Core Team 2024) to implement linear mixed effects models in *lme4* (v1.1.35.5; Bates et al., 2015) and *lmerTest* (v3.1.3; Kuznetsova et al., 2017). We used *tidyverse* and *ggplot2* to process data and create plots (v2.0.0; Wickham et al., 2019; v3.5.1; Wickham 2016).

We first modeled leaf temperature as a function of fixed effects of leaf conductance, measurement day (and thus weather, Table 1), leaf shape, and a leaf shape*measurement day interaction; we included sampling quadrant as a random effect. The second model had the same structure except leaf conductance was the response and leaf temperature was a fixed predictor, to account for the relationship between temperature and conductance. In this second model, the random effect of sampling quadrat was estimated to be zero, so we dropped this term and used a linear model.

We used *emmeans* (v1.10.5; Lenth 2024) to obtain marginal mean leaf temperatures and stomatal conductances from each model, grouped by leaf shape and measurement day. We compared extracted means of leaf shape and day using Tukey’s Post Hoc tests (Figure 1c and 1d).

Residuals of the model for stomatal conductance measures were normally distributed; residuals from models of temperature were not. We used *bestNormalize* (v1.9.1; Peterson 2021) which identified an Ordered Quantile (ORQ) normalization as the best transformation of temperature. Analysis of ORQ-transformed, log-transformed, and square-root transformed data yielded identical results in direction and statistical significance. We present analyses and means with temperature data in original units for ease of interpretation.

## Acknowledgements

We want to thank Radana Molnarova, Julio de Almeida, and all other staff at the Koffler Scientific Reserve for accommodating and supporting this project. We are incredibly grateful to the University of Toronto’s Centre for Global Change Science for their support and funding (YKS). We thank Amanda Peake for providing feedback on the manuscript and assistance in the field, and NSERC Canada for funding in the form of Discovery Grants (JRS) and CGS M and PGS D graduate fellowships (JAB).

## Author Contributions

Yash Kumar Singhal: conceptualization, data curation, formal analysis, investigation, methodology, writing -original draft, writing -review editing, visualization.

Julia Anne Boyle: conceptualization, funding acquisition, supervision, writing -original draft, writing -review editing, formal analysis.

John R. Stinchcombe: conceptualization, funding acquisition, methodology, project administration, resources, supervision, writing -review editing, writing -original draft, formal analysis.

